# Role of RgsA in oxidative stress resistance in *Pseudomonas aeruginosa*

**DOI:** 10.1101/2020.03.03.974683

**Authors:** Shuyi Hou, Jiaqing Zhang, Xiaobo Ma, Qiang Hong, Lili Fang, Gangsen Zheng, Jiaming Huang, Yingchun Gao, Qiaoli Xu, Xingguo Zhuang, Xiuyu Song

## Abstract

*Pseudomonas aeruginosa* is an extremely common opportunistic pathogen in clinical practice. Patients with metabolic disorders, hematologic diseases, malignancies, who have undergone surgery or who have received certain treatments are susceptible to this bacterium. In addition, *P. aeruginosa* is a multidrug-resistant that tends to form biofilms and is refractory to treatment. Small regulatory RNAs are RNA molecules that are 40–500 nucleotides long, possess regulatory function, are ubiquitous in bacteria, and are also known as small RNA (sRNA). sRNAs play important regulatory roles in various vital life processes in diverse bacteria and their quantity and diversity of regulatory functions exceeds that of proteins. In this study, we showed that deletion of the sRNA RgsA decreases the growth rate and ability to resist different concentrations and durations of peroxide in *P. aeruginosa*. These decreases occur not only in the planktonic state, but also in the biofilm state. Finally, protein mass spectrometry was employed to understand changes in the entire protein spectrum. The results presented herein provide a description of the role of RgsA in the life activities of *P. aeruginosa* at the molecular, phenotypic, and protein levels.

## Introduction

*Pseudomonas aeruginosa* is ubiquitous in nature and the most common opportunistic pathogen. Patients with metabolic disorders, hematologic diseases, malignancies, who have undergone surgery, or who have received certain treatments are susceptible to this organism. In addition, *P. aeruginosa* is multidrug-resistant. *P. aeruginosa* has low pathogenicity, but exhibits extremely strong resistance against complex environments, particularly chemicals and drugs. Therefore, it often causes persistent infection, leading to protracted disease. *P. aeruginosa* tends to form biofilms, and 65% of chronic diseases in humans are related to biofilms. Even though there are no genetic changes in bacteria in biofilms, they exhibit a different phenotype known as the biofilm phenotype. An important characteristic of this phenotype is strong resistance to continuously changing environments. Investigations of the life processes of *P. aeruginosa* to understand its growth, pathogenicity, and resistance characteristics, as well as to identify genes and regulation involved in these phenotypes and potential new therapeutic targets are currently topics in *P. aeruginosa* research.

Small regulatory RNAs (sRNAs) are small RNA molecules that are 40–500 nucleotides long, possess regulatory function, and are ubiquitous in bacteria. sRNAs play important regulatory roles in various vital life processes in diverse bacteria^[1]^. Generally, the expression of sRNA molecules is induced by certain transcription factors under specific environmental cues, metabolic signals, or survival stresses. sRNAs can regulate one to many targets or act as global regulatory factors to participate in many biological processes. sRNA regulation is usually conserved, and the quantity and diversity of its regulatory effects exceed that of proteins.

RgsA, which was discovered from a whole genome search conducted in 2008, is a sRNA that is present in *P. aeruginosa* and *P. fluorescens*. The predicted gene length of RgsA is 120 bp, it is directly regulated by the σ factor RpoS and indirectly regulated by the GacS/GacA two-component system. Northern blot analysis showed that RgsA exists as two transcripts during the logarithmic phase and one transcript during the stationary phase. RgsA deletion causes hydrogen peroxide resistance in bacteria to decrease, but the specific mechanism responsible for this is unknown^[2]^. RNA expression differences between the planktonic and biofilm states of *P. aeruginosa* have been studied. The RgsA expression level in biofilms at 48 h of culture was 1000 times that of planktonic bacteria at 4 h (exponential phase) and 12 h (stationary phase), suggesting that RgsA plays an important role in the high resistance of bacterial biofilms^[3]^.

In this study, we studied the effects of RgsA deletion on oxidative stress resistance and survival stress in *P. aeruginosa* to provide a basis for identifying a complete effector mechanism for RgsA.

## Materials and methods

### Bacteria, plasmid, and growth conditions

Table 1 shows information on bacterial strains and plasmids used in this experiment. The culture media used in this experiment consisted of Luria broth (LB) and nutrient agar plates. For screening, 100 μg/ml ampicillin, 25 μg/ml (*E. coli*) and 100 μg/ml (*P. aeruginosa*) tetracycline, 10 μg/ml (*E. coli*) and 40 μg/ml (*P. aeruginosa*) gentamycin, and 100 μg/ml (*E. coli*) and 300 μg/ml (*P. aeruginosa*) spectinomycin were used. *E. coli* and *P. aeruginosa* were inoculated from glycerol stock onto nutrient agar plates and cultured overnight, after which a single colony was selected and subcultured in LB, then cultured to an optical density at 600 nm of 0.5 (OD_600_≈0.5) which was the pre-logarithmic growth phase. The bacterial suspension obtained was centrifuged and washed before experiments were conducted.

**Table 1.** Bacterial strains and plasmids.

| Strain of plasmid | Characteristics | Reference or source |
| --- | --- | --- |
| ***E.coli*** |  |  |
| DH5α | supE44ΔlacU169 (ϕ80lacZΔM15) hsdR17 recA1 end A1 gyrA96 thi-1 relA1 | Solarbio |
| SM10 | thi thr leu tonA lacy supE recA::RP4-2-Tc::Mu:: | [4] |
| ***P. aeruginosa*** |  |  |
| PAO1 | wild type | [5] |
| PAO1 (pJNT) | PAO1 contain plasmids pJNT Gen<sup>r</sup> Tet<sup>r</sup> | This study |
| PAO1ΔRgsA | ΔrgsA:: aacC1 Gen<sup>r</sup> | This study |
| PAO1ΔRgsA (pJR1) | PAO1ΔRgsA contain plasmids pJR1 Gen<sup>r</sup> Tet<sup>r</sup> | This study |
| PAO1ΔRgsA(pJNT) | PAO1ΔRgsA contain plasmids pJNT Gen<sup>r</sup> Tet<sup>r</sup> | This study |
| **Plasmids** |  |  |
| pUCGM | aacC1 source, Ap<sup>r</sup> Gm<sup>r</sup> | [6] |
| pMD 19-T | Ap<sup>r</sup> | Takara |
| pEX18-Ap | Ap<sup>r</sup>, oriT<sup>+</sup> sacB<sup>+</sup>, gene replacement vector with MSC from pUC18 | [7] |
| pEX18-TE | Tet<sup>r</sup>, oriT<sup>+</sup> sacB<sup>+</sup>, gene replacement vector with MSC from pUC18 | This study |
| pJN105 | araC-pBAD cloned in pBBR1 MCS-5; Gm<sup>r</sup> | [8] |
| pJNT | The plasmid pJN105 contains Tet gene cassette, Gen<sup>r</sup> Tet<sup>r</sup> | This study |
| pJR1 | RgsA cassette in pJNT, Gen<sup>r</sup> Tet<sup>r</sup> | This study |

### DNA and RNA extraction

Bacterial genomic DNA was extracted using a QIAquick PCR Purification Kit (Qiagen, Germany). A kit from Tiangen Biotech (China) was used for plasmid DNA extraction. A QIAquick Gel Extraction Kit (Qiagen) was used for plasmid recovery. The Trizol method was used for RNA extraction^[9]^.

### Determination of the transcription start site of RgsA

The 5’RACE method was employed to determine the transcription start site of RgsA. Briefly, *P. aeruginosa* PAO1 was cultured in LB culture medium until OD_600_≈0.5 (the pre-logarithmic phase), at which time total RNA was extracted. A cDNA reverse transcription system was used to synthesize the first cDNA strand. After purification and recovery, TdT and dATP were used to add a poly(A) tail to the 5’ terminus of the cDNA. The product of this reaction was used as a template. The RACE-anchor-T17 anchor primer containing 17–20 Ts at the 3’ end as well as the specific primers RgsA-R3 and GM-PA2958-R1 were used for three rounds of nested PCR to amplify the transcripts. After recovery of the DNA fragments, a T vector was constructed by ligation with the Pmd 19-T Vector. This vector was transformed into DH5α competent cells. Next, 10 tubes of positive strains were used for plasmid extraction, of which plasmids from six tubes were used for sequencing and alignment with the gene library to determine the transcription start site.

### Construction of RgsA-deletion strain

The PAO1 overnight culture was collected and genomic DNA was extracted. Two pairs of primers (Hind III-PA2957-F1/PA2959-PstI-PA2958-R1 and PA2959-PstI-PA2958-F1/Bam HI-PA2960-R3) were used for PCR amplification of the upstream homology arm PA2958 and the downstream homology arm PA2959 in the RgsA gene. The primers were designed so that the ends of the products PA2958 and PA2959 contained 47 bp complementary sequences. The mixture of PA2958 and PA2959 was subjected to overlap PCR with the primer pair Hind III-PA2957-F1/BamH I-PA2960-R3.After gel purification and digestion with restriction enzymes (Hind III and BamH I), the PCR product *PA2958-PA2959* gene fragment was cloned into the suicide vector pEX18Ap. The ligated vector was subsequently transformed into DH5α competent cells, which were screened on ampicillin plates. Plasmids were extracted and digested using *Pst* I, after which they were ligated with the gentamicin resistance gene fragment that was obtained from *Pst* I digestion of the pUGM plasmid. The ligated product was then transformed into competent *E. coli* SM10 to construct the recombinant *E. coli* SM10 strain (pEX18-*PA2958-PA2959*-*aacC1*), which was screened from ampicillin + gentamicin plates. Two-parent hybridization was used to transfer the recombinant plasmid in *E. coli* SM10 (pEX18-*PA2958-PA2959*-*aacC1*) to PAO1. The suicide vector-mediated homologous recombination method was used and SPE+GN was employed for screening to obtain the PAO1ΔRgsA mutant. DNA was extracted from the screened mutants for validation.

### Construction of RgsA-overexpression strain

According to the 5’RACE results, EcoRI-rgsA-F1/PstI-rgsA-R1 was used to amplify the RgsA short transcript fragment. NcoI was used to digest pJN105, which was ligated with the *tet* gene that was obtained from NcoI digestion of pEX18-TE to obtain the pJNT plasmid containing tetracycline resistance. The pJNT plasmid and *rgsA* short transcript gene fragment were double digested using *Eco*R I and *Pst*I followed by ligation to obtain the recombinant plasmid pJR1, which was subsequently transformed to obtain the *E. coli* SM10 (pJR1) recombinant strain. This was followed by screening on TET+GN plates. Finally, two-parent hybridization was used to transfer the recombinant plasmid in *E. coli* SM10 (pJR1) to PAO1 ΔRgsA to obtain the PAO1ΔRgsA (pJR1) overexpression strain.

### Construction of parallel experimental strain

The same method was used to construct the parallel experimental strain PAO1 (pJNT) using the pJNT plasmid and PAO1, PAO1ΔRgsA (pJNT) using the pJNT plasmid and PAO1ΔRgsA.

### Growth curve

A total of 0.5 ml of PAO1 and PAO1ΔRgsA overnight culture was subcultured into 50 ml of LB culture medium and cultured at 37°C and 200 rpm. The OD_600_ was measured every 15 minutes from the early logarithmic phase to the mid-logarithmic phase. Each measurement was conducted in triplicate and means were calculated. The OD_600_ was measured every 30 minutes from the late logarithmic phase to the stationary phase. Each measurement was carried out in triplicate and means were calculated, after which the values were recorded and used to plot a curve.

### Plate survival experiment

PAO1 (pJNT), PAO1ΔRgsA (pJNT) and PAO1ΔRgsA (pJR1) were separately diluted with 0.85% Nacl+0.1% L-Arabinose solution to OD_600_≈0.3 before 10-fold serial dilution using the same solution was conducted to generate six concentrations. Next, the diluted samples were inoculated on 0.7 mM CHP, 0.7 mM TBH, and 2 mM H_2_O_2_ plates and cultured for 24 hours, after which bacterial survival was observed.

### MTT assay for quantification of cell viability

PAO1 (pJNT), PAO1ΔRgsA (pJNT) and PAO1ΔRgsA (pJR1) were resuspended in 0.85% Nacl+0.1%L-Arabinose solutions. The concentrations of the suspensions were adjusted to an OD_600_ of 0.15–0.17, after which they were aliquoted into EP tubes of 1 ml each. The following oxidative stress simulation conditions were then generated: 40 mM H_2_O_2_ for 30 minutes, 10 mM H_2_O_2_ for 1 hour, and 10 mM for 5 hours. Every sample was repeated three times. After the reaction was completed, the solutions were centrifuged and the pellets were washed. Next, 100 μl 0.85% NaCl solution was used to resuspend the pellet and 10 μl MTT (3-(4,5-Dimethylthiazol-2-yl)-2,5-diphenyltetrazolium bromide) was added to each 100 μl resuspended solution. The samples were then incubated at 37°C for 60 minutes with the caps opened. Next, the solutions were centrifuged at 12,000 rpm for 1 minute, after which 99 μl of supernatant was aspirated and 1250 μl DMSO solution was used to dissolve the precipitate. Finally, the OD550 was measured after 10–15 minutes of reaction.

### In vitro biofilm culture

A flow apparatus was used for biofilm culture. PAO1 (pJNT), PAO1ΔRgsA (pJNT) and PAO1ΔRgsA (pJR1) were separately diluted with 0.85% Nacl+0.1% L-Arabinose solution to a McFarland standard of 0.5. Acid-treated glass slides were then immersed in the prepared bacterial suspension and incubated at room temperature for 24 hours. Next, the slides were placed in a flow apparatus. The perfusate was 10% LB+0.1% L-Arabinose solution, which was subjected to continuous culture for 72 hours at a flow rate of 0.05 ml/s. PBS was then added as the perfusate and planktonic bacteria on the glass slides was flushed for 10 minutes at a flow rate of 0.5 ml/s. The perfusion device was then removed and the glass slides were taken out and washed once with PBS. Next, the biofilm slides obtained in the previous step were immersed in 10 mM, 20 mM, or 30 mM H_2_O_2_ solutions for 30 minutes. PBS was subsequently used to repeatedly wash the glass slides, which were slightly dried, then covered with SYTO9/PI dye that had been diluted 1000 times. Staining was conducted for 20 minutes, after which the glass slides were removed from the solution, excess water was removed and the slides were kept moist until observed using a fluorescence microscope.

### Detection of protein expression differences by two-dimensional electrophoresis

PAO1 and PAO1ΔRgsA were added to 0.5 ml lysis-buffer and sonicated for lysis. After high speed and low temperature centrifugation, the supernatant was collected and 1.5 ml 20% TCA/acetone was added and mixed evenly. The solution was then allowed to stand at 4°C for 30 minutes, after which it was subjected to high speed centrifugation. Next, the precipitate was washed three times with 10% acetone containing 20% mm DTT, after which it was subjected to high speed and low temperature centrifugation. The pellet was then collected and air-dried before being re-dissolved in 50 μl swelling solution. Two-dimensional electrophoresis was then conducted using Ettan IPGphor 3 for isoelectric focusing, Ettan Dalt6 for parallel electrophoresis, and ImageMaster 2D Platinum for image analysis (all from GE Healthcare) according to the manufacturer’s instructions, after which the gel was silver-stained and protein differences on the gel were compared. Differentially expressed proteins were extracted and sent to BGI Group (Shenzhen) for mass spectrometry identification.

## Result

### RgsA transcription start site

Based on the sequencing and sequence alignment results, there are two RgsA transcripts in *P. aeruginosa* that are 90 bp and 300 bp in length. The transcription start sites of the short and long transcripts were 86 bp and 157 bp downstream of the start site of the NCBI annotated gene fragment, respectively, and a common transcription terminator may be present (Fig 1).

**Fig 1.**
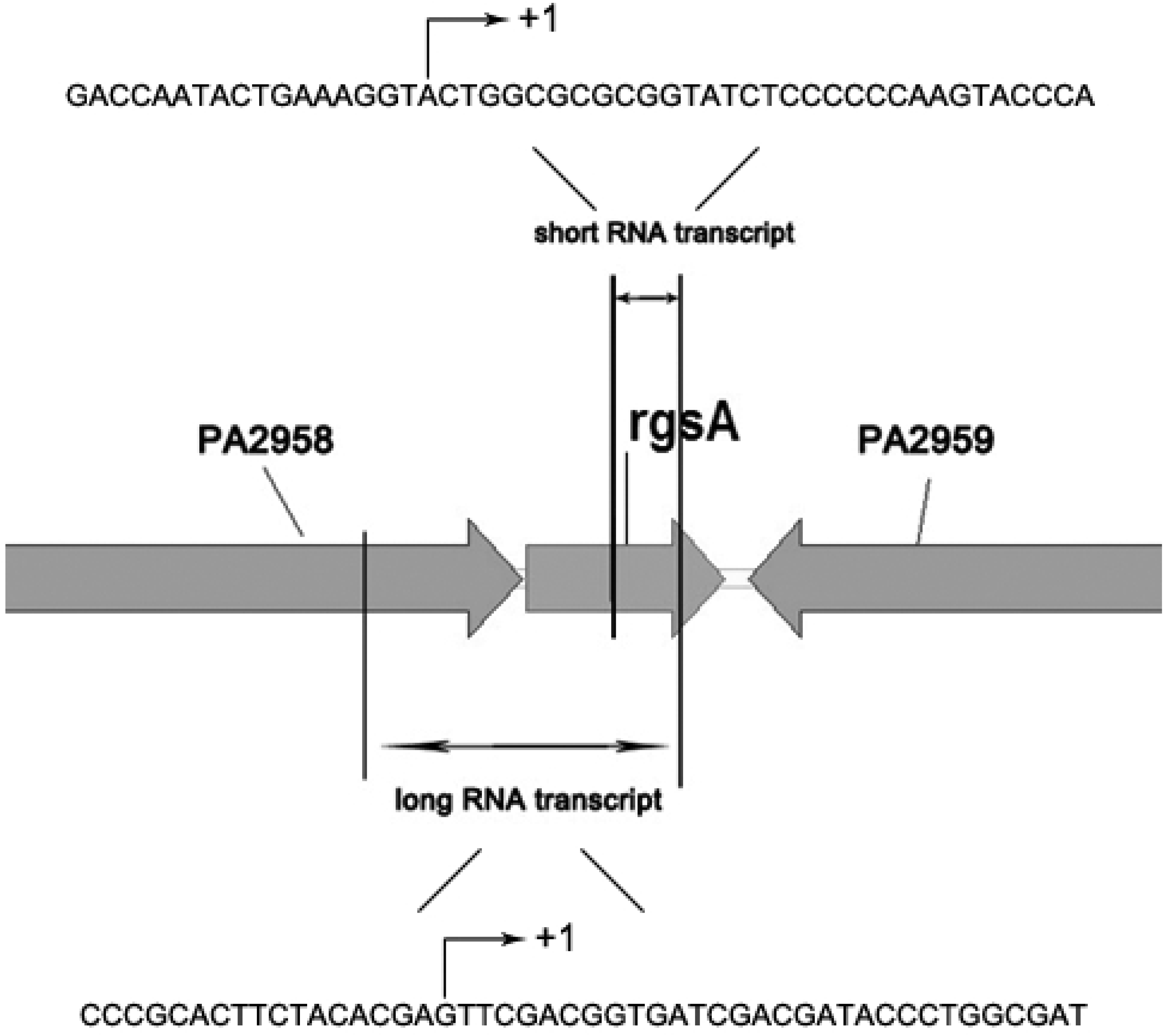
Schematic diagram of the RgsA transcript site and sequence.

### Growth curve

In a previous experiment, we noticed that the turbidity of the RgsA-deletion strain after overnight culture was lower than that of the wild-type strain. Therefore, we measured the OD_600_ values of PAO1 and PAO1ΔRgsA for 10 hours to determine if RgsA will affect the growth rate of *P. aeruginosa*. The difference in growth rate between the RgsA deletion mutant and the wild-type strain could be as high as 8 times at the start of the logarithmic phase. When the same starting inoculum was used for culture, the total bacterial concentration was 8 fold less than that of the wild-type strain after the stationary phase was reached (Fig 2).

**Figure 2.**
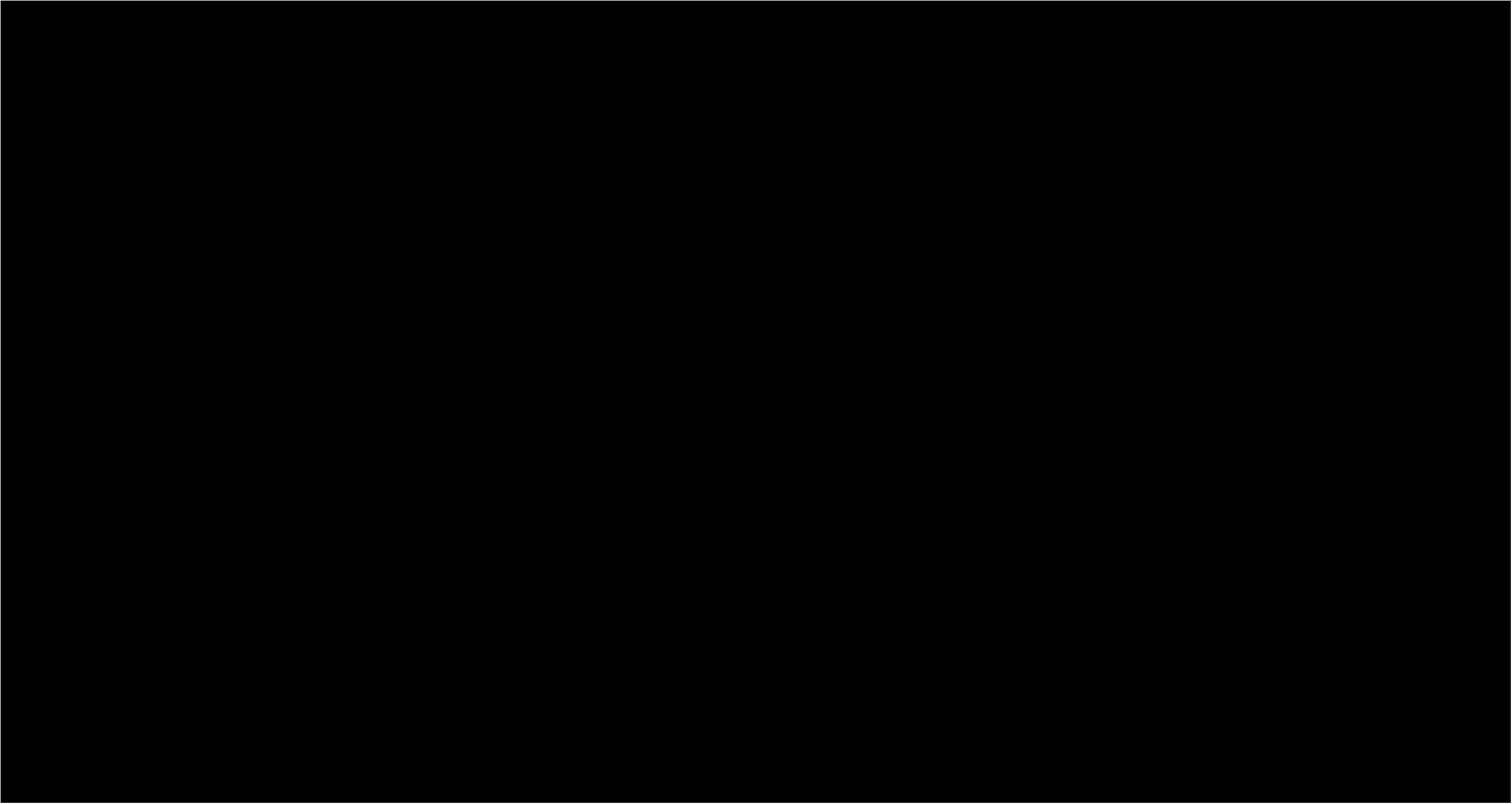
Growth curve of PAO1 and PAO1ΔRgsA in LB culture medium.

### Oxidative stress resistance

To investigate the effects of different oxidative stress substances on RgsA mutant growth, we conducted plate survival experiments to measure the resistance of PAO1ΔRgsA(pJNT), PAO1ΔRgsA (pJR1), and PAO1 (pJNT) to hydrogen peroxide and organic peroxides. Hydrogen peroxide and organic peroxide resistance in the RgsA deletion mutation were significantly decreased. After introduction of the expression vector into the mutant and induction with 0.1% L-Arabinose, oxidative stress resistance basically recovered to levels near the wild-type strain (Fig 3).

**Figure 3.**
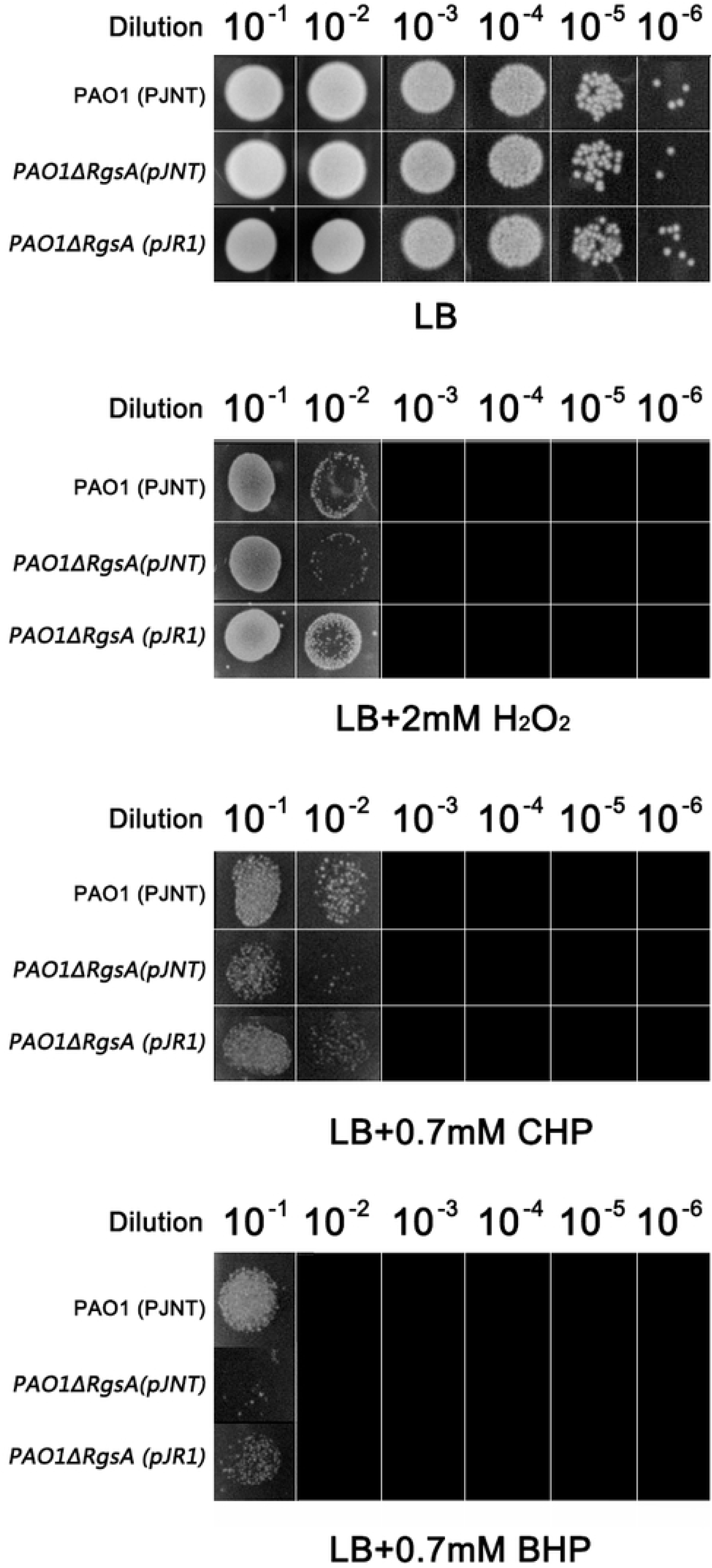
plate survival experiments of PAO1ΔRgsA(pJNT), PAO1ΔRgsA (pJR1), and PAO1 (pJNT)

To study the effects of oxidative stress concentration and duration on RgsA mutant growth, we employed an MTT assay and designed two low concentration (10 mM) and long duration (2 h, 5 h), and one high concentration (40 mM) and short duration (30 minute) hydrogen peroxide stimulation experiments to test the three strains. We found that, as duration and concentration increased, the number of surviving RgsA deletion bacteria was reduced compared with the wild-type strain. Under short duration and high concentration stress conditions, the overexpression strain did not recover to similar resistance as the wild-type strain (Table 2, Fig 4).

**Table 2.** PAO1ΔRgsA(pJNT), PAO1ΔRgsA (pJR1), and PAO1 (pJNT) MTT measurement results.

|  | PAO1 (pJNT) | PAO1ΔRgsA(pJNT) | PAO1ΔRgsA (pJR1) |
| --- | --- | --- | --- |
| 40 mM H<sub>2</sub>O<sub>2</sub> 30 min | 0.710621 | 0.491116 | 0.559638 |
| 10 mM H<sub>2</sub>O<sub>2</sub> 2 h | 0.719343 | 0.699317 | 0.768371 |
| 10 mM H<sub>2</sub>O<sub>2</sub> 5 h | 0.817538 | 0.647131 | 0.789501 |

**Figure 4.**
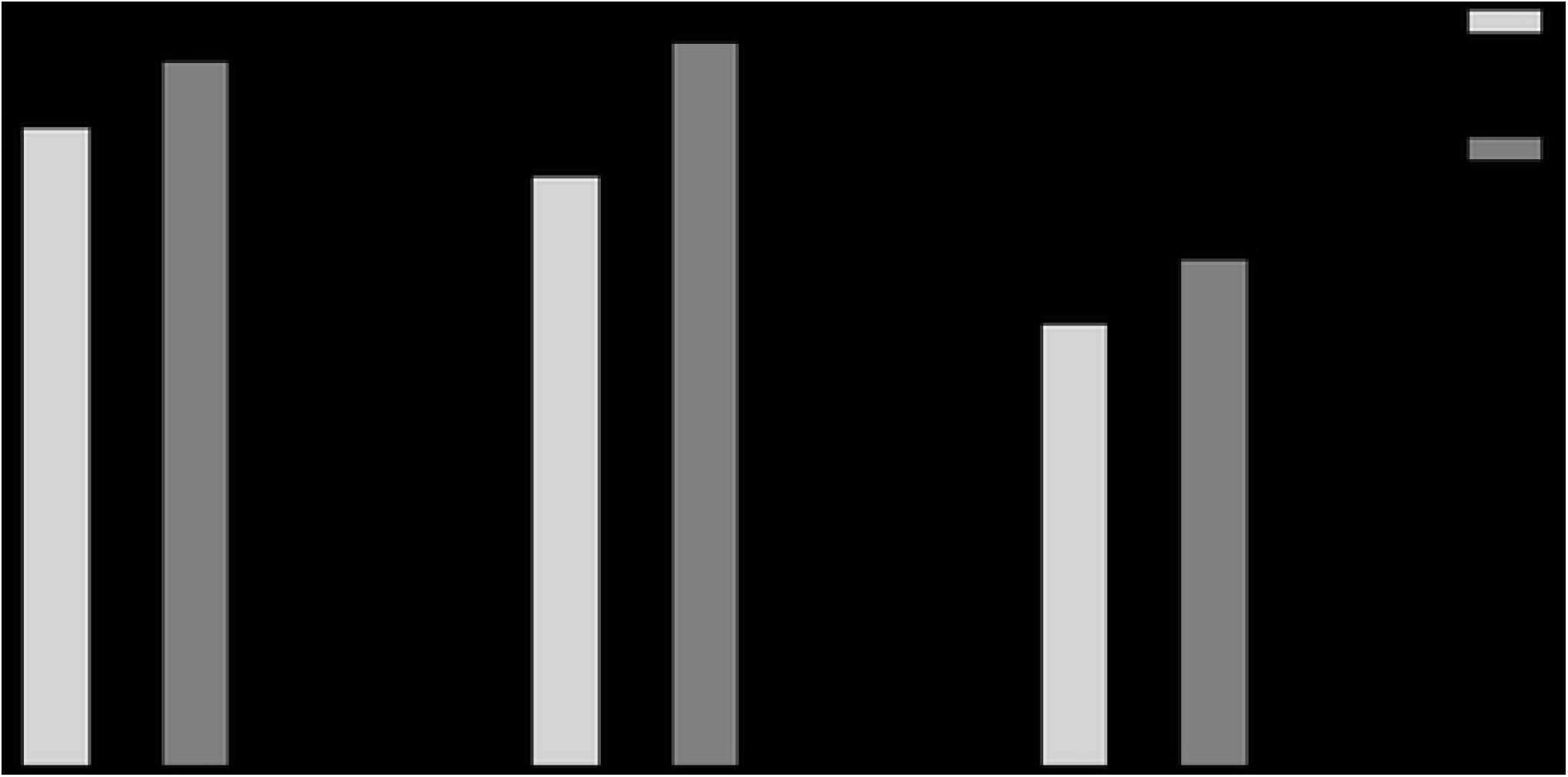
PAO1ΔRgsA(pJNT), PAO1ΔRgsA (pJR1), and PAO1 (pJNT) MTT measurement changes.

### Oxidative stress resistance in the biofilm state

We used a flow apparatus to form biofilms after 3 days of culture, then added different concentrations of hydrogen peroxide for *in vitro* simulation of oxidative stress, after which SYTO9/PI staining followed by fluorescence microscopy was conducted. When the hydrogen peroxide concentration was 20 mM, more dead bacteria appeared in the RgsA deletion mutant treatment, and this difference became greater when 30 mM hydrogen peroxide as used. However, the overexpression strain could recover to levels close to those of the wild-type strain, which is consistent with oxidative stress responses in the planktonic state (Fig 5).

**Figure 5.**
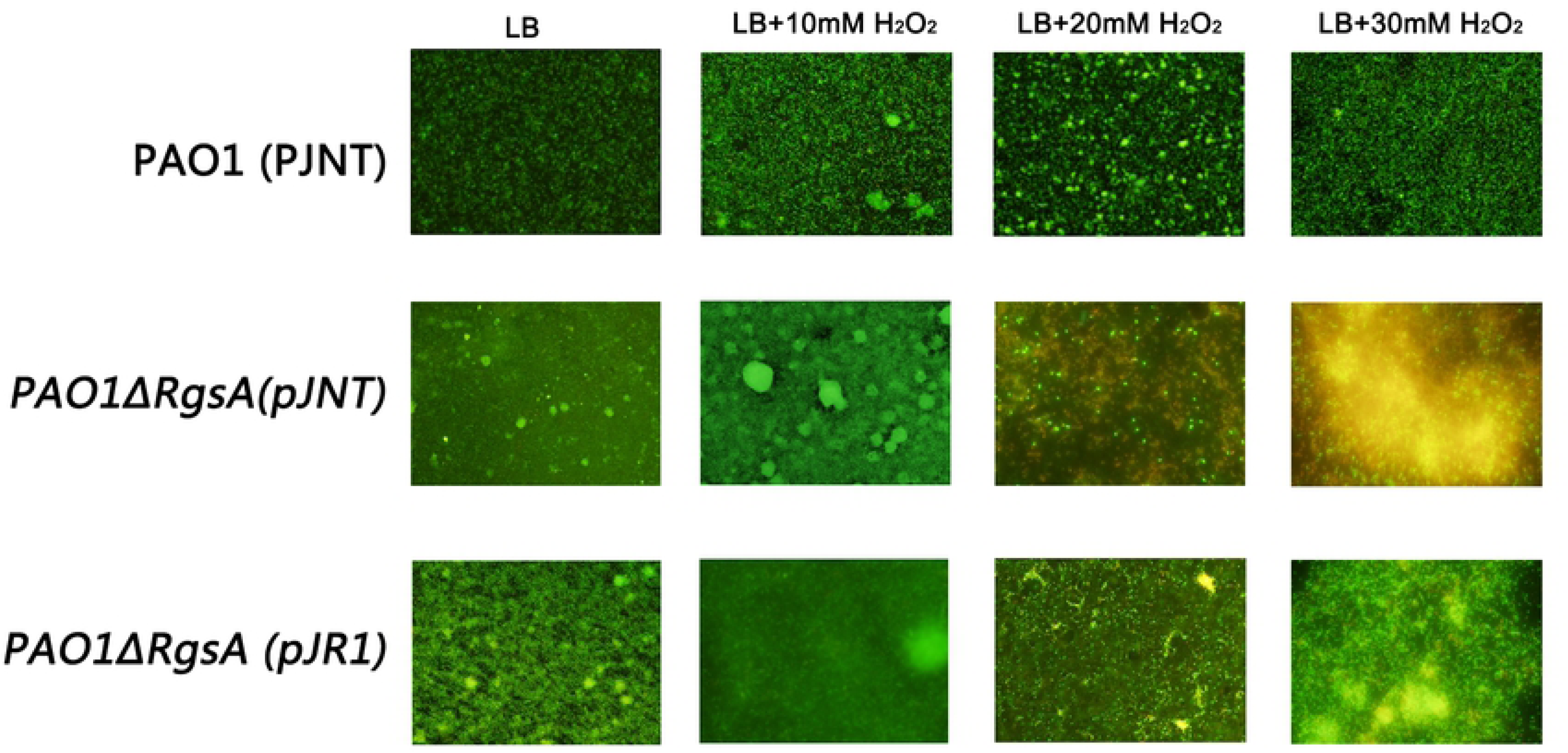
Oxidative stress resistance in the biofilm state.

### Detection of protein expression differences by two-dimensional electrophoresis

To understand the effects of RgsA on global protein expression in *P. aeruginosa*, we selected PAO1 and PAO1ΔRgsA for two-dimensional electrophoresis. On the protein gel, we identified 18 spots with significant differences in expression, of which 16 proteins were downregulated and two were upregulated. We conducted mass spectrometry identification analysis based on the PAO1 protein library, as well as other *P. aeruginosa* protein libraries for some proteins that were not found in the PAO1 protein library. Detailed protein information is shown in Table 3.

**Table 3.** List of differentially expressed proteins identified upon two-dimensional electrophoresis of PAO1 and PAO1ΔRgsA.

| Index | Protein ID | Regulation | Protein mass | Isoelectric Point | Coverage rate | Description |
| --- | --- | --- | --- | --- | --- | --- |
| 1 | PA39016_002180023 | down | 53373.4<br>3 | 5.88 | 42.86% | probable outer membrane protein precursor [*Pseudomonas aeruginosa* PAO1 PAO1-UW] |
| 2 | PSPA7_5705 | down | 53022.3<br>1 | 5.58 | 47.19% | membrane efflux protein [*Pseudomonas aeruginosa* PA7] |
| 3 | PALES_40651 | down | 48023.2 | 8.69 | 50.81% | alkaline protease secretion protein AprE [*Pseudomonas aeruginosa* PAO1 PAO1-UW] |
| 4 | PSPA7_2715 | down | 46799.6<br>7 | 9.96 | 38.32% | membrane protein CzcC [*Pseudomonas aeruginosa* PA7] |
| 5 | PA39016_002180023 | down | 53373.4<br>3 | 5.88 | 33.75% | probable outer membrane protein precursor [*Pseudomonas aeruginosa* PAO1 PAO1-UW] |
| 6 | PA39016_002330023 | down | 47153.7<br>8 | 6.82 | 37.14% | transcription termination factor Rho [*Pseudomonas aeruginosa* PAO1 PAO1-UW] |
| 7 | PALES_10551 | down | 51198 | 8.95 | 34.87% | conserved hypothetical protein [*Pseudomonas aeruginosa* PAO1 PAO1-UW] |
| 8 | NCGM2_5074 | down | 24134.3 | 7.6 | 44.95% | protein [*Pseudomonas aeruginosa* NCGM2.S1] |
| 9 | PADK2_07705 | up | 30911.2<br>9 | 9.38 | 27.37% | putative ABC-type phosphate/phosphonate transport system, periplasmic component [*Pseudomonas aeruginosa* M18] |
| 10 | PA2G_02188 | down | 33904.5<br>1 | 5.7 | 31.08% | putative ATPase [*Pseudomonas aeruginosa* M18] |
| 11 | PALES_38391 | up | 33147.1<br>9 | 8.6 | 35.64% | probable binding protein component of ABC transporter [*Pseudomonas aeruginosa* PAO1 PAO1-UW] |
| 12 | PA39016<br>_001710<br>046 | down | 19042.8<br>8 | 9.37 | 37.16% | conserved hypothetical protein<br>[*Pseudomonas aeruginosa* PAO1<br>PAO1-UW] |
| 13 | PALES_<br>54511 | down | 15003 | 9.77 | 42.45% | conserved hypothetical protein<br>[*Pseudomonas aeruginosa* PAO1<br>PAO1-UW] |
| 14 | PSPA7_5<br>705 | down | 53022.3<br>1 | 5.58 | 44.28% | membrane efflux protein<br>[*Pseudomonas aeruginosa* PA7] |
| 15 | PA39016<br>_002180<br>023 | down | 53373.4<br>3 | 5.88 | 38.51% | probable outer membrane protein<br>precursor [ *Pseudomonas*<br>*aeruginosa* PAO1 PAO1-UW] |
| 16 | PALES_<br>10551 | down | 51198 | 8.95 | 34.43% | conserved hypothetical protein<br>[*Pseudomonas aeruginosa* PAO1<br>PAO1-UW] |
| 17 | PA39016<br>_002030<br>004 | down | 17561.5<br>3 | 4.77 | 68.46% | suppressor protein DksA<br>[*Pseudomonas aeruginosa* PAO1<br>PAO1-UW] |
| 18 | PA5247 | down | 18049.2 | 7.93 | 45.34% | hypothetical protein<br>[*Pseudomonas aeruginosa* PAO1<br>PAO1-UW] |

## Discussion

Aerobic and facultative anaerobic bacteria primarily employ aerobic respiration for energy metabolism. During this process, aberrant electron transport on the electron transport chain or cell oxidoreductase may produce reactive oxygen intermediates (ROIs) such as superoxide (O^2-^), hydrogen peroxide (H_2_O_2_), and hydroxyl free radicals (·OH). In addition, bacteria may be exposed to exogenous ROIs, particularly during infection of the human body, when human phagocytes (such as neutrophils) produce significant oxygen-dependent antibacterial effects. These ROIs may cause oxidative damage to cellular proteins, cell membranes, and genetic substances such as DNA. Bacteria possess a series of protective systems to resist oxidative stress, which involve antioxidant proteins (superoxide dismutase, catalase, and peroxidase), iron sequestration, free radical scavengers, DNA-binding proteins, and DNA repair enzymes.

In this study, we first determined the length of the RgsA transcripts and their transcription start sites. The results of northern blot analysis showed that RgsA may be present as two transcripts with different lengths during the logarithmic phase, while the 300 bp transcript is undetectable during the stationary phase^[2]^. The total RNA samples used in this study were obtained from *P. aeruginosa* in the pre-logarithmic phase (OD_600_≈0.5), and there were two types of transcripts with different lengths present in the RNA samples. It was difficult to obtain the full-length 5’RACE, and the PCR-recovered fragments consisted of a mixture of different start sites that varied by several to dozens of bases that were indistinguishable, therefore, the sequences with different length were then aligned with the *P. aeruginosa* RgsA gene sequence in the NCBI database and the result with the longest alignment was considered to the be the start site of the 5’ end of the transcript. Based on the sequencing and sequence alignment results, there are two RgsA transcripts in *P. aeruginosa* with lengths of 90 bp and 300 bp, which corresponded to the results of Northern blot analysis. The transcription direction was the same and common transcription stop sites may be present. These findings provided a basis for subsequent construction of effective deletion and overexpression strains.

Our study focused on the role of RgsA in oxidative stress resistance. RgsA deletion resulted in decreased resistance to hydrogen peroxide and organic peroxide by *P. aeruginosa* in the planktonic and biofilm states, and this decrease was directly proportional to an increase in the concentration of hydrogen peroxide and duration of the treatment. Decreased tolerance to the external environment may have influenced the growth rate and final concentration. In biofilm experiments, we also found that the RgsA deletion strains formed thinner biofilms that tended to detach from carrier surfaces. However, the biofilms recovered to levels near those of the wild-type strain after introduction of the overexpression plasmid.

Finally, we conducted mass spectrometry analysis of the protein expression spectrum of the deletion and wild-type strains. Comparison of these spectra clearly showed that RgsA knockout significantly downregulated protein expression, including various outer membrane protein and efflux pump proteins, which are vital for bacterial adaptation and stability in a variable environment. In addition, some differentially expressed proteins participated in regulation of transcription, secretion and virulence in *P. aeruginosa*. Among these proteins, the DksA protein is a novel regulatory factor and its downregulation may result in incomplete DNA replication or asynchronous DNA replication and slowed cell growth^[10]^. Additionally, DksA participates in the posttranscriptional regulation of extracellular virulence factors in *P. aeruginosa*, can affect the expression of rhamnolipid and LasB elastase^[11]^, and forms an alkaline phosphatase with the membrane fusion protein AprE, ABC transporter protein AprD, and outer membrane protein AprF of the OprM family, which is a type 1 secretion system. Reduced DksA expression may decrease the extracellular secretion of alkaline protease AprA and AprX, thereby affecting *P. aeruginosa* infection, particularly its pathogenicity during early infection^[12]^.

Based on analysis of RgsA from the DNA structure to its biological function and protein expression, we understand how sRNA regulates RNA. Although RgsA itself does not have functions of its own, it participates in various regulatory responses in bacteria, thereby affecting bacterial survival, metabolism, and pathogenicity. With the development of machine learning and various bioinformatics techniques, more sRNA and their targets will be identified and validated, which is vital to obtaining a complete understanding of bacteria.

## Acknowledgments

We would like to thank Professor Bin Huang of the First Affiliated Hospital, Sun Yat-sen University for the Pseudomonas aeruginosa PAO1, Professor Yihe Ge of Ludong Universtiy, Professor Kangmin Duan of Northwest University, Professor Ningyi Zhou of Wuhan Institute of Virology, Professor Jianqiang Lin of Shangdong University, Professor Rui Zhou of Huazhong Agricultural University for the plasmids and strains.

